# Genome of tropical bed bug *Cimex hemipterus* (Cimicidae, Hemiptera) reveals tetraspanin expanded in bed bug ancestor

**DOI:** 10.1101/2024.04.25.591052

**Authors:** Sean Tsz Sum Law, Wenyan Nong, Chade Li, Tze Kiu Chong, Ho Yin Yip, Thomas Swale, Siu Wai Chiu, Roger Yat-Nork Chung, Hon-Ming Lam, Samuel Y.S. Wong, Hung Wong, Jerome H.L. Hui

## Abstract

*Cimex* species are ectoparasites that exclusively feed on warm-blooded animals such as birds and mammals. Three cimicid species are known to be persistent pests for humans, including the tropical bed bug *C. hemipterus*, common bed bug *C. lectularius*, and Eastern bat bug *Leptocimex boueti*. To date, genomic information is restricted to the common bed bug *C. lectularius*, which limits understand their biology and to provide controls of bed bug infestations. Here, a chromosomal-level genome assembly of *C. hemipterus* (495 Mb) containing on 16 pseudochromosomes (scaffold N50 = 34 Mb), together with 9 messenger RNA and small RNA transcriptomes were obtained. Comparison between hemipteran genomes, we found that the tetraspanin superfamily were expanded in the *Cimex* ancestor. This study provides the first genome assembly for the tropical bed bug *C. hemipterus*, and offers an unprecedented opportunity to address to questions relating to bed bug infestations, as well as genomic evolution to hemipterans more widely.

## Introduction

Cimicids in the Hemiptera are insects that feed on hosts’ blood. Despite the family Cimicidae contains 91 described species and can be divided into 6 subfamilies and 23 genera (Krinsky 2002), most people refer bed bugs to the three better known species where humans serves as their hosts - tropical bed bug *C. hemipterus* (Fabricius 1803), common bed bug *C. lectularius* (Linnaeus 1758), and Eastern bat bug *Leptocimex boueti* (Brumpt 1910). People bitten by bed bugs could have reactions ranging from localised urticaria, bullous reactions, to anaphylaxis in rare cases (Thomas et al 2004). With globalisation happened in the last half a century together with the easiness of cimicids, bed bug is now considered as worldwide resurging problem with *C. hemipterus* and *C. lectularius* being the major species implicated in human infestations (CDC 2022).

*Cimex hemipterus* and *C. lectularius,* similar to other hemipterans or true bugs, are hemimetabolous insects and its life cycle includes eggs, nymphal instars, and adults. All their instars and adults require to take blood meals from humans, although other mammals and birds can also be their hosts in the absence of humans. Female bed bugs can lay 5 eggs a day and eggs will further hatch into first instar nymphs in 4 to 12 days. A blood meal is essential for molting to the next developmental stage. Copulation or traumatic insemination happened between adults to fertilise eggs to complete the life cycle. As adults generally live 6 to 12 months and can survive for long periods of time without feeding, together with the occurrence of insecticide resistance, bed bug control and eradication is challenging and considered as a global problem (Goddard and deShazo 2009; Romero et al 2007).

Obtaining the genome sequence of animals are useful for obtaining deeper understanding of their biology, as well as providing a valuable resource for varieties of follow-up studies. In the case of bed bugs, understanding the underlying molecular mechanisms of their mating, feeding, and digestion on human blood, could potentially be useful to make new measures for further control and eradication (Reinhardt et al 2007). Nevertheless, as of to date, only the genome of common bed bug *C. lectularius* has been sequenced and analysed (Rosenfeld et al 2016). Here, we aim to obtain a high-quality genome and transcriptomic resources of the missing key infestation bed bug for human, tropical bed bug *C. hemipterus*, to shed light on the situation.

## Methodology

### Sample collection, Genome sequencing and assembly

bed bugs were collected in different districts in Hong Kong including Fanling, Kwai Hing, Kwun Tong, Shatin, Tsing Yi and Tsuen Wan. Genomic DNA was extracted with QIAamp DNA Mini Kit (Qiagen), and sent to BGI Hong Kong Company Limited for stLFR sequencing on DNBSEQ G400 to obtain 2×100bp paired-end reads (Wang et al 2019; Peters et al 2019). Dovetail Omni-C library was further prepared as described (Lieberman-Aiden et al 2009), and sequenced on Illumina HiSeq X to give 2 × 150 bp paired-end reads. The stLFR reads were used for de novo assembly according to the stLFR pipeline (https://github.com/BGI-Qingdao/stlfr2supernova_pipeline)(Weisenfeld et al 2017; Wang et al 2019), with default parameters. The pseudohap style output dedupe contigs and Dovetail Omni-C library reads were used as input data for HiRise, a software pipeline specifically designed to use proximity ligation data to scaffold genome assemblies (Putnam et al. 2016). Sequences from the Dovetail Omni-C library were aligned to the draft input assembly using bwa (https://github.com/lh3/bwa). The separation of Dovetail Omni-C read pairs mapped within the draft scaffolds was analysed using HiRise to build a likelihood model for the genomic distance between read pairs. The model was used to identify and break putative misjoins, score potential joins and make joins above a threshold. ). Estimation of genome size, repeat content and heterozygosity was carried out by GenomeScope2 with K-mer at 31 (Vurture et al 2017), which was then compared with two *Cimex lectularius* genomes in NCBI (Aseesion numbers GCA_000648675.3 and GCA_001460545.1). Syntenic analysis between the X ,Y chromosome of the Heteroptera genomes (*Aelia acuminata*, GCA_911387785.2; *Acanthosoma haemorrhoidale* GCA_930367205.1; *Aradus depressus* GCA_963662175.1) and the *C. hemipterus* genome was run using SyMAP (Soderlund et al 2011). Details of the sequencing data can be found in Supplementary Table S1.

### Transcriptome sequencing

Total RNA was extracted from adult samples using the mirVana miRNA Isolation Kit (Ambion) according to the manufacturer’s protocol. The quality of extracted total RNA was checked using a Nanodrop spectrophotometer (Thermo Scientific), gel electrophoresis and an Agilent 2100 bioanalyser (Agilent RNA 6000 Nano Kit). Qualified samples were sent to Novogene (HK) Company Limited for library construction and sequencing. PolyA-selected RNA sequencing libraries were prepared using the TruSeq RNA Sample Prep Kit v2. Insert sizes and library concentrations of the final libraries were examined using an Agilent 2100 Bioanalyser instrument (Agilent DNA 1000 Reagents). Similarly, isolated small RNA was submitted to Novogene (HK) Company Limited for HiSeq small RNA library construction and 50 bp single end (SE) sequencing. Details of the sequencing data can be found in Supplementary Table S1. Raw sequencing reads from five transcriptomes were pre-processed with quality trimmed by trimmomatic (v0.39 with parameters ”ILLUMINACLIP:TruSeq3-PE.fa:2:30:10 SLIDINGWINDOW:4:5 LEADING:5 TRAILING:5 MINLEN:25”) (Bolger et al 2014) and contamination removed by Kraken2 (database version k2_standard_20210517) (Wood et al 2019). The processed reads were used for the following gene model prediction.

### Gene model prediction

The gene model was predicted as previously described (Li et al. 2020). Briefly, gene models were trained and predicted using funannotate (v1.8.9, https://github.com/nextgenusfs/funannotate) with the parameters “--repeats2evm -- protein_evidence uniprot_sprot.fasta --genemark_mode ET --busco_seed_species fly -- optimize_augustus --busco_db insecta --organism other --max_intronlen 350000”. Gene models from several prediction sources, including GeneMark, Augustus high-quality predictions (HiQ), pasa, Augustus, GlimmerHM and snap, were passed to Evidence Modeler and used to generate gene model annotation files and applied in PASA to update EVM consensus predictions, add UTR annotations and models for alternatively spliced isoforms. Protein-coding gene models were then blasted (blastp) to the NCBI nr and swissprot databases using diamond (v0.9.24), specifying the following parameters: --more-sensitive -- evalue 1e-3, and mapped with transcriptome reads using HISAT2 (version 2.1.0) (Kim et al. 2019). Gene models with no homology to known proteins in the nr and swissprot databases, and without mRNA support, were removed from the final version. Gene model expression profiles were generated from different individuals by extracting transcript per million (TPM) values from mapped reads using StringTie (PERTEA et al. 2015).

### Gene orthology and gene family evolution analysis

Orthologous groups were inferred from the longest representative protein sequences of 29 Hemipterans including *C. hemipterus* downloaded from NCBI (https://www.ncbi.nlm.nih.gov/datasets/genome/GCF_000648675.2/) and 21 arthropod taxa using OrthoFinder v2.5.2 (Emms and Kelly 2019) (Supplementary Table S5). The default settings were used and the multiple sequence alignments option “-MSA” was activated to generate a maximum-likelihood of species tree, which was further inferred for divergence times with r8s (Sanderson 2003) and TimeTree web database (http://timetree.org/). The orthologous groups and species tree were used as input in CAFE5 analysis (Mendes et al 2020). Candidates of expanded gene families were inspected by protein family search with HMMER (version 3.3.1; cut-off E-value <10−5) (Eddy 2011), from which each gene candidate was validated with conserved Domain Database (CDD) via RPS-BLAST (Yang et al 2020). Subsequently, gene families of interest were investigated for sequence alignment using MAFFT v7.455 (Katoh and Standley, 2013), followed by phylogenetic analysis with FastTree (Price et al 2010) and visualization in evolview v3 (Subramanian et al 2019). For the classification of tetraspanin families, reference sequences from Huang et al (2005) were used. The pairwise sequence identity was computed by Clustal Omega (v1.2.4, Sievers et al 2011). Microsynteny between *C. hemipterus* and *C. lectularius* was identified with MCScanX (Wang et al 2012) followed by visualization with *gggenomes* (Hackl et al 2021). DeepTMHMM (Hallgren et al 2022) was employed to annotate the transmembrane regions of tetraspanin.

### Gene family annotation and gene tree building

*Drosophila melanogaster* Hox and ParaHox genes were retrieved from HomeoDB (Zhong & Holland, 2011) to search for Hox and ParaHox genes in both *C. hemipterus* and *C. lectularius* using tBLASTn (Altschul et al., 1997) with threshold of E-value of <10^-3^. Reference homeobox gene sequences were retrieved from HomeoDB and a published study (Qu et al., 2020). For hormonal and anticoagulation genes annotation, amino sequences were first obtained from KEGG or NBCI gene database. The obtained sequences were then used to retrieve the corresponding genes from *C. hemipterus* genome using the using tBLASTn with threshold of E-value of <10^-3^. All annotated genes were further proceeded to phylogenetic analyses using Neighbor-Joining method with MEGA7 (Kumar et al., 2016) and Maximum likelihood method using IQTREE (version 2.2.0.3) (Schrempf et al., 2016). The phylogenetic tree was then visualized with iTOL (Letunic & Bork, 2021). Expression of the annotated hormonal genes were visualized by heatmap illustration using TBtools (Version No.1.112) (Chen et al 2020).

### MicroRNA annotations

Quality of small RNA sequencing reads were checked using FastQC (Andrews et al 2010), and reads between 18 and 27 bp in length were then mapped to the genome using mapper.pl. MicroRNAs were predicted using miRDeep2.pl from miRDeep2 (Mackowiak et al 2011). Expression profiles and quantification were generated by the miRDeep2 script quantifier.pl with 0 mismatches in read-to-predecessor mapping and other default parameters. The raw count table was used for differential expression analysis by Trinity’s run_DE_analysis.pl and analyse_diff_expr.pl scripts using the edgeR method, where at least two replicates had calculated counts per million (CPM) values ≥ 1, and cut-offs were set at 0.05 p-values (FDR) and 2-fold changes. Predicted microRNAs in *C. hemipterus* were manually checked by using BLASTn in miRBase (Kozomara et al 2019) with default settings, as well as aligning them with reference microRNA hairpin sequences from MirGeneDB (Fromm et al 2022). The genomic locations of all annotated microRNAs were visualised with Tbtools (Chen et al 2020), and the distance between neighboring members of a microRNA cluster is set to 10 kb (Kozomara et al 2019). The microRNA of *Cimex lectularius* was retrieved from InsectBase 2.0 (Mei et al 2022). The presence of non-annotated miRNAs hairpin of *Cimex lectularius* was also checked by using BLASTn to the genomes with miRNA hairpin of *C. hemipterus* as reference.

### Repeat annotation

Transposable elements (Tes) were annotated as previously described (Baril et al 2022) using the automated Earl Grey TE annotation pipeline (version 1.2, https://github.com/TobyBaril/EarlGrey). In brief, this pipeline first involved the identification of known Tes from the lepidopteran subset of Dfam (release 3.4) and RepBase (release 20,181,026) (Hubley et al 2016). An automated ‘BLAST, Extract, Extend’ process was then used to identify de novo Tes and extend consensus boundaries (Platt et al 2016). Redundant sequences were removed from the consensus library before the genome assembly was annotated with the combination of known and *de novo* TE libraries. The annotations were then processed to remove overlaps and to defragment the annotations prior to the final TE quantification.

## Results and Discussion

### A high-quality genome of *C. hemipterus*

Here we present a high-quality chromosome-level genome assembly of *C. hemipterus* (2n = 32), with 97.25 % of the genome sequence contained on 16 pseudomolecules (∼495.4 Mb, scaffold N50 = 34.6 Mb) (Figure 1B, 1C; Supplementary Table S2), from which macrosynteny revealed Chr 2 and Chr 3 are X_1_ and X_2_ choromosomes that correspond to the male karyotype (2n = 28 + X_1_X_2_Y) (Sadílek et al 2019) (Supplementary Figure 1), where the Y chromosome was not assmebled in the genome. The size of the assembled genome is comparable with the estimated size of the genome (484.2 Mb) and the two *C. lecutlarius* genomes (553 Mb and 517 Mb for GCF_000648675.2 and GCA_001460545.1, respectively) (Supplementary Figure 2). A total of 33,254 predicted gene models were annotated using the transcriptomes generated in this study, including 16,500 protein-coding genes and 16,754 tRNA genes, and the genome completeness estimated by BUSCO analyses was 96.40 % (Figure 1C; Supplementary Table S3). We have also identified a total of 22.81 % repeat contents in the *C. hemipterus* genome (Figure 1D), with many of these newly discovered repetitive elements are relatively understudied in current databases (unclassified: 8.98%). Among the remaining identifiable repeats, LINE elements and DNA transposons are the most abundant (LINEs: 7.21%, DNA transposons: 3.82%), whereas SINEs, LTR elements and rolling circle elements are only present in low proportions (LTR elements: 0.43%, rolling circle elements: 0.07%, SINEs: 1.16%) (Figure 1D), which is similar to the bed bug *C. lectularius* (Rosenfeld et al. 2016). Our genome in this study provides the first chromosome-level genome in the Cimicidae (Supplementary Table S4).

**Figure 1.**
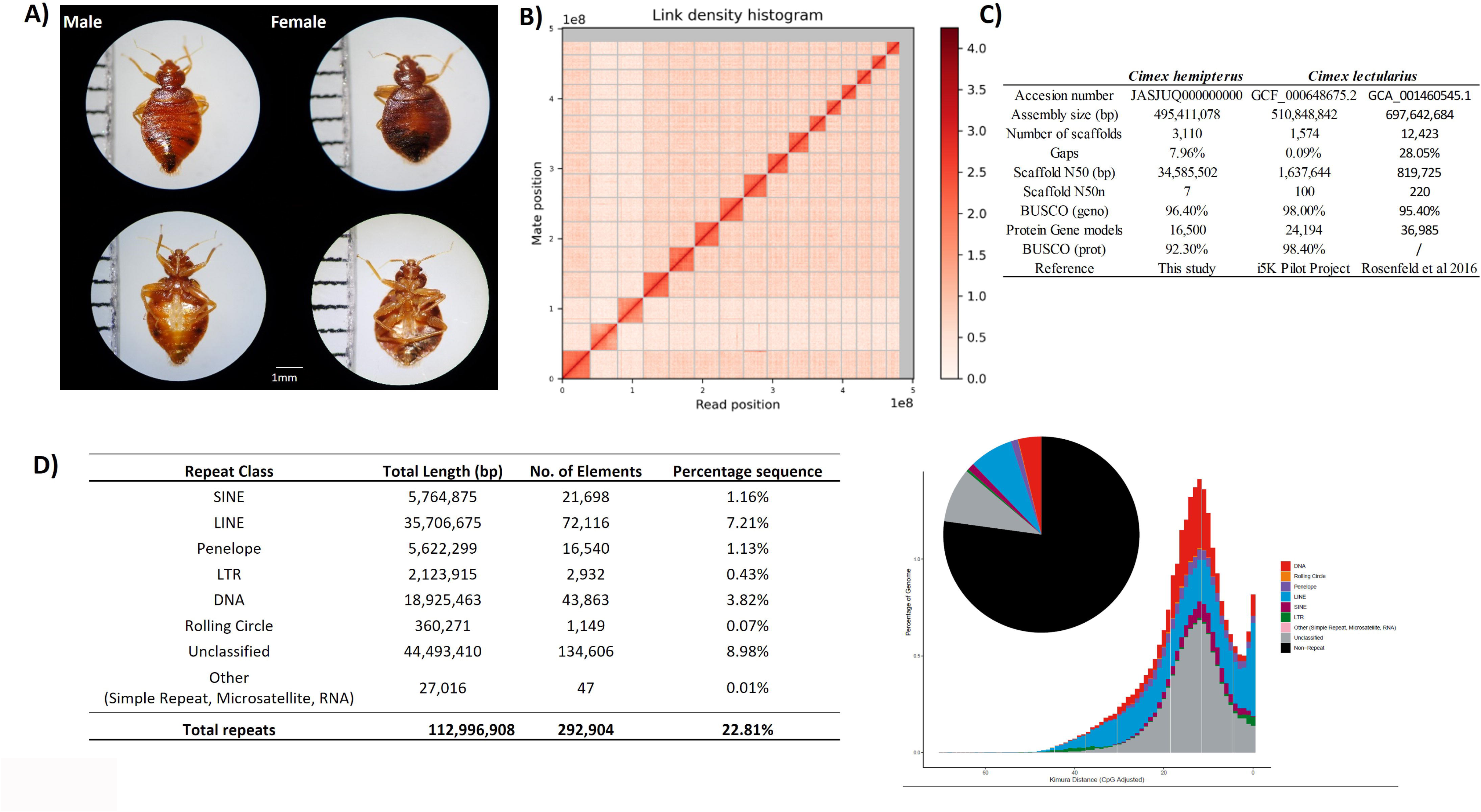
A) Photographs of male and female of tropical bed bug *Cimex hemipterus*. B) Hi-C interaction of 16 linkage groups; C) Statistics of *C. hemipterus* and *C. lectularius* genome assemblies; D) Transposable elements in *C. hemipterus*.

### Anticoagulant proteins and gene family evolution in the *Cimex* lineage

Haematophagous animals such as the bed bugs have the abilities to secrete anticoagulants to prevent blood clotting and triggering other immune response of the hosts in order to obtain blood meals. Similar to *C. lectularius*, we can also identify genes involved in the production of different anticoagulant proteins (Supplementary Figure 3-5, Supplementary Table S12, S13).

To understand what other adaptation could have arisen in the *Cimex* ancestor, genome-wide gene gain and loss analyses across insects with the inclusion of this new genomic resource were carried out (Figure 2A and B). We have found that two transmembrane protein families, FAR-17a/AIG1-like protein and tetraspanin (also known as TM4SF), were significantly expanded in both *Cimex* genomes (Figure 2C). A larger number of gene copies of FAR-17a/AIG1-like proteins was detected in *C. hemipterus* (n = 31) when compared to *C. lectularius* (n = 14). The difference was contributed by tandem gene duplicates in *C. hemipterus*, including a cluster of 9 FAR-17a/AIG1-like protein coding genes on chromosome 16 (Supplementary Figure 6A). Although all annotated FAR-17a/AIG1-like proteins were expressed in *C. hemipterus* (Supplementary Figure 6B), little is known about their function in insects as FAR-17a/AIG1-like proteins were mainly characterized in vertebrate models as an androgen-responsive gene and a role player in cancer-related processes (Huang et al 2021). On the other hand, in addition to known insect tetraspanin orthologous families (Huang et al 2005), the expansion in tetraspanin was contributed by a newly identified family termed “N-3”, which was also found in hemipteran *Orius laevigatus* (Supplementary Figure 7 and 8). Moreover, not only some tetraspanins in the N-3 cluster contained a novel “CCS” cysteine pattern of large extracellular domain (EC2) instead of the conserved “CCG” signature, but most N-3 tetraspanins were expressed in male only (Supplementary Figure 8). It is noteworthy that tandem gene duplications of N-3 tetraspanins were also identified in both *Cimex* species, in which larger gene clusters were observed in *C. hemipterus* and led to larger gene number of tetraspanins than in *C. lectularius* (Supplementary Figure 8A). In animals, tetraspanin are well known membrane-spanning proteins that function as molecular scaffolds contribute to development, reproduction and immunity (Termini and Gillette 2017). In fruit flies *Drosophila*, species-specific tetraspanin *Sunglasses* (DmTsp42Ej) is required for retina development, and for flies with extended light exposure but having reduced *Sunglasses* expression resulted the inability to regulate rhodopsin (Xu et al 2004). Given that bed bugs are nocturnal and are able to detect colours and objects at very low background light conditions (Singh et al 2015), whether *Cimex* tetraspanins are involved in visual detection would be an interesting story to test. Understanding the functions of these *Cimex-*specific tetraspanin and whether they can be used as candidates for combating bed bugs, similar to the development of potential therapeutic targets in human diseases, warrants further investigation.

**Figure 2.**
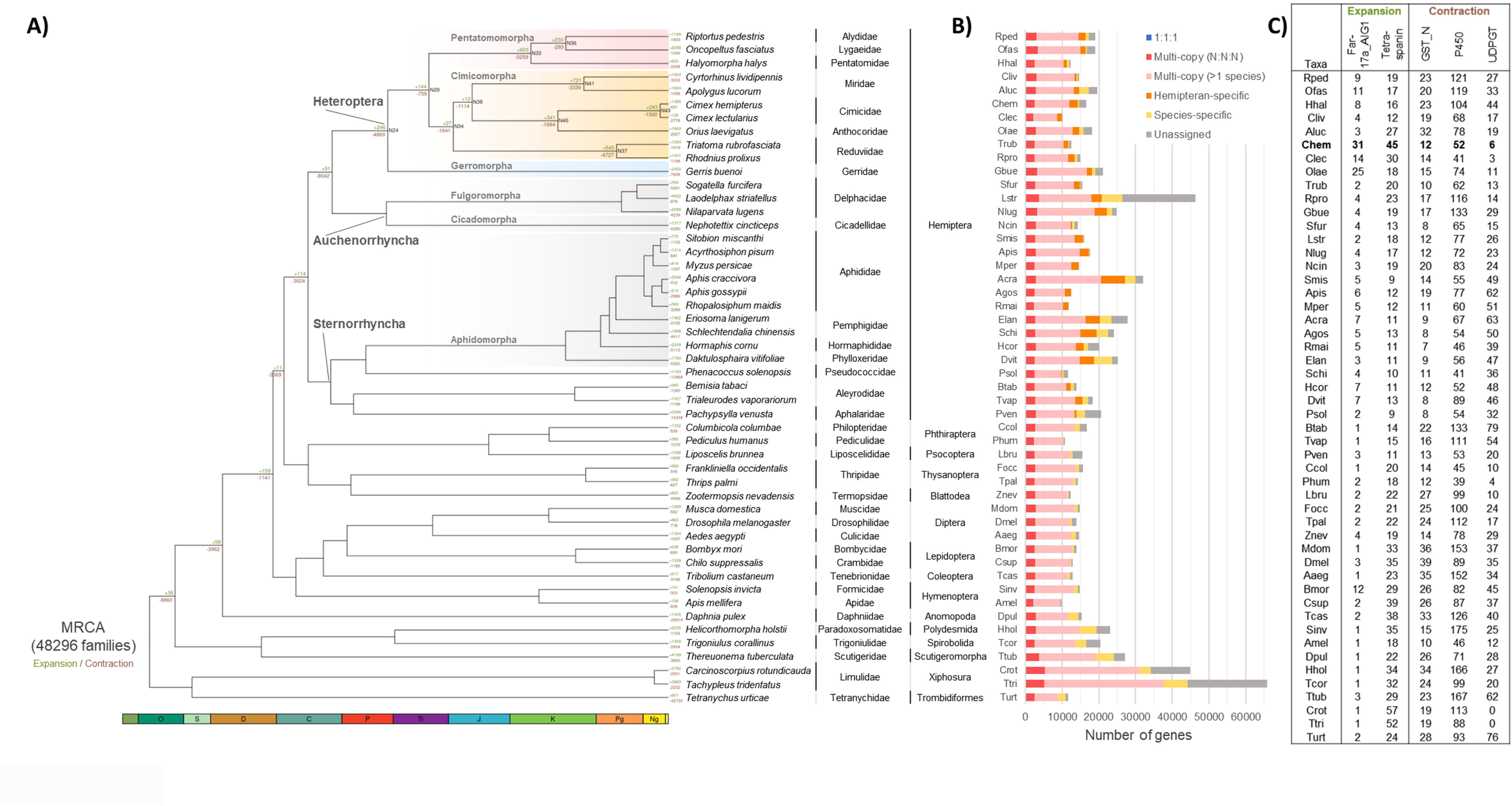
A) Summary of gene family expansion and contraction in *Cimex* and other reference animals. MRCA, most recent common ancestor; B) Number and distribution of genes assigned to different orthologous groups; C) Gene count table with expanded or contracted protein family domains.

### Developmental and hormonal genes in *C. hemipterus*

Homeobox genes such as Hox and ParaHox genes are transcription factors involved in regulating animal development, and its genomic organisation can help understanding how animals evolved (e.g. Nong et al 2020). A total of 13 Hox and ParaHox genes were identified in each of *C. hemipterus* and *C. lectularius* genomes (Figure 3A, Supplementary Table S6, Supplementary Figure 9). In *C. hemipterus* genome, the majority of Hox genes are clustered on the same scaffold with exception of one posterior Hox gene, while the two ParaHox genes are located on two different scaffolds. The situation contrasts that of the *C. lectularius*, where majority of Hox genes are located on two different scaffolds, presumably due to lower scaffold N50 (Figure 3A). It is worth noting that the posterior Hox gene were both located on a different scaffold to majority of Hox genes in both *Cimex* genomes, suggesting the translocation event could be dated back to the *Cimex* last common ancestor. The Hox and ParaHox genes expressed largely similarly between sexes (Figure 3B, Supplementary Table S7).

**Figure 3.**
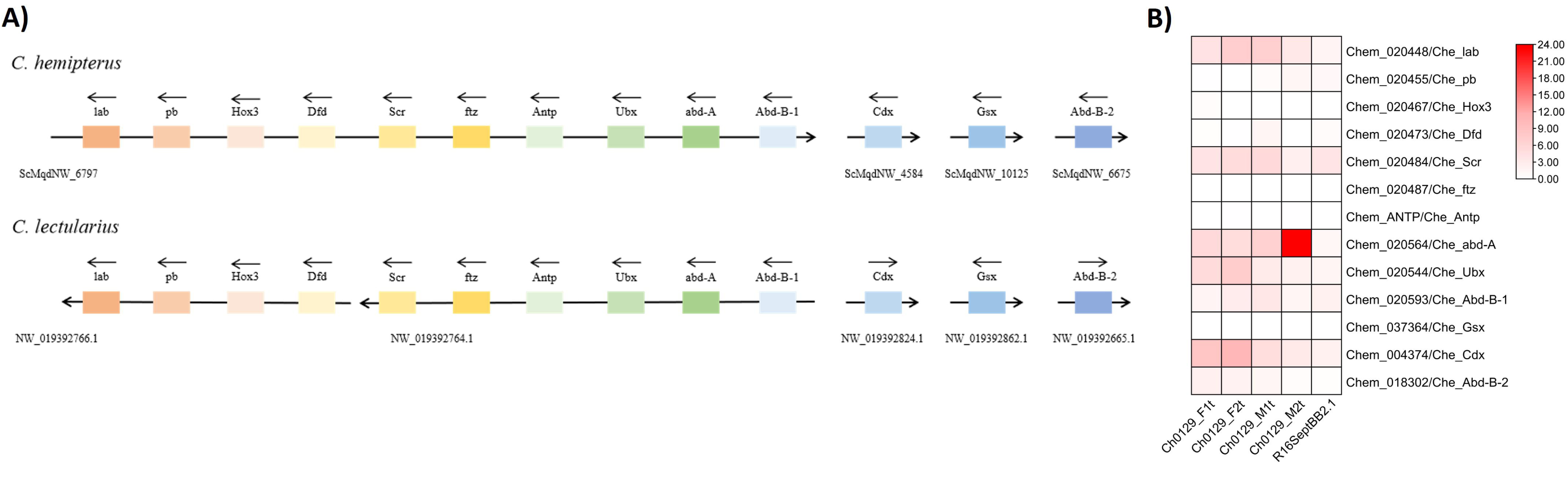
A) Genomic locations of Hox and ParaHox genes in *C. hemipterus* and *C. lectularius*. B) Expression of Hox and ParaHox genes in *C. hemipterus*.

Sesquiterpenoid hormones such as juvenile hormones control development and reproduction in insects, and are potential targets for controlling insect growth and numbers. In the *C. hemipterus* genome, genes involved in the MVA pathway, including *ACAT, HMGCS, HMGCR, MVK, PMVK, DPMD, and FPPS* were identified (Figure 4A; Supplementary Table S8; Supplementary Figure 10-16), suggesting *C. hemipterus* starts the sesquiterpenoid production by mevalonate (MVA) pathway (Tobe and Bendena 1999; Bendena et al 2018; Tsang et al 2020; So et al 2022). However, *FPPP* and *FOHSDR* that code for enzymes to convert farnesyl-PP to farnesol and farnesal respectively as in many other arthropods were not detected in the *C. hemipterus* genome (Baker et al 1983; Sperry & Sen 2001; Mayoral et al 2009; Nyati et al 2013; Satvaveanthan et al 2021). Instead, gene members involved in the isoprenylation pathway, namely *FNTB*, *STE24*, *ICMT*, and *PCYOX1*, were detected, indicating that *C. hemipterus* may produce farnesal through isoprenylation pathway (Zhang & Casey 1996) (Supplementary Figure 17-20). In addition, we could identify orthologues of *ALDH III*, *JHAMT*, and *CYP15A1* (Figure 4A; Supplementary Table S8; Supplementary Figure 21-23), suggesting FA, MF and JHSB3 were being synthesized in *C. hemipterus*, which is consistent with other Hemipterans (Rivera-Perez et al 2003; Kotaki et al 2009; Tsang et al 2020; Villalobos-Sambucaro et al 2020; So et al 2022). Comparing the gene expression level between the two sexes, we found that there was a higher gene expression level of *HMGCR* and lower gene expression level of a *STE24* copy in the two female samples (Ch0129_F1t, Ch0129_F2t) than the two male samples (Ch0129_Mt1, Ch0129_Mt2). However, the gene expression level of sequiterpenoid genes of another bed bug species *C. lectularius* do not show any sex-biased expression (Supplementary Table S9; Supplementary Figure 24).

**Figure 4.**
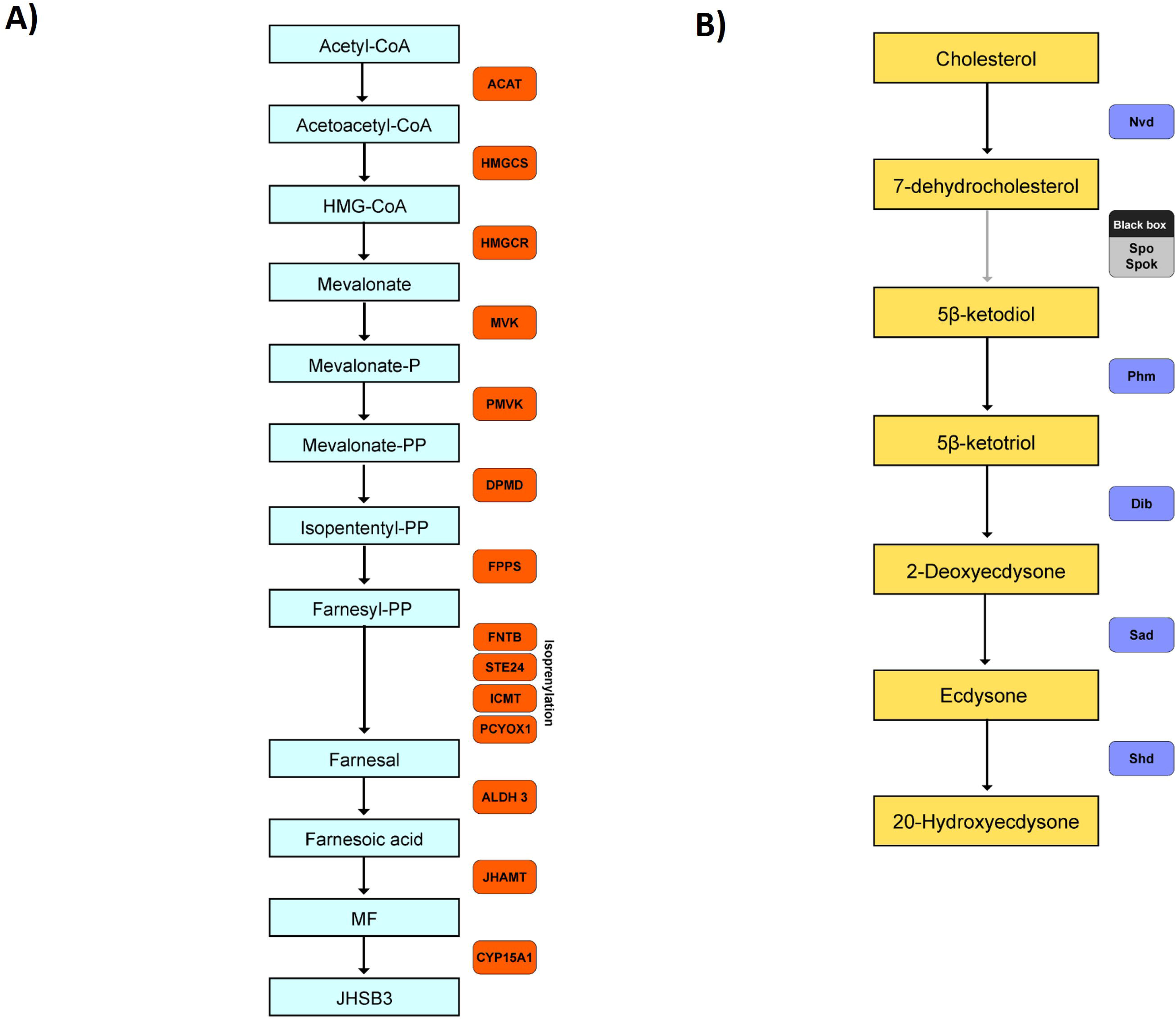
Sesquiterpenoid and Ecdysteroid pathway genes in tropical bed bug *Cimex hemipterus*. A) Schematic diagram of sesquiterpenoid biosynthetic pathway genes identified in *C. hemipterus*. *ACAT*, Acetyl-CoA Acetyltransferase; *HMGCS*, hydroxymethylglutaryl-CoA synthase; *HMGCR*, 3-hydroxy-3-methylglutaryl-CoA reductase; *MVK*, mevalonate kinase; *PMVK*, phosphomevalonate kinase; *DPMD*, diphosphomevalonate decarboxylase; *FPPS*, farnesyl diphosphate synthase; *FNTB,* protein farnesyl transferase; *STE24*, endopeptidase; *ICMT*, protein-S-isoprenylcysteine O-methyltransferase; *PCYOX1*, prenylcysteine oxidase; *ALDH 3*, aldehyde dehydrogenase 3; *JHAMT*, juvenile hormone acid O-methyltransferase. B) Sesquiterpenoid pathway genes identified in *C. hemipterus*. *Nvd*, Neverland; *Spo*, Spook; *Spok*, Spookier; *Phm*, Phantom; *Dib*, disembodied; *Sad*, Shadow; *Shd*, Shade.

Other than sesquiterpenoids, ecdysteroids are another group of important hormones that regulate growth (such as molting and metamorphosis) and sexual maturation of insects (Cheong et al 2015). In the *C. hemipterus* genome, *Spo*/*Spok*, *Nvd, Phm, Dib, Sad,* and *Shd* could all be identified (Figure 4B; Supplementary Table S10; Supplementary Figure 25), suggesting *C. hemipterus* shared a similar ecdysteroid biosynthesis pathway to other insects, which cholesterol is converted to 7-dehydrocholesterol (7dC) and then to 5β-ketodiol (Namiki et al 2005; Niwa and Niwa. 2014; Ono et al 2006; Magico and King-Jones 2011; Niwa et al 2010; Warren et al 2009), followed by further hydroxylation where 5β-ketotriol, 2-deoxyecdysone, ecdysone, and 20-Hydroxyecdysone are synthesized respectively (Niwa and Niwa 2014). Comparing the gene expression level between the two sexes, except for *Nvd*, the expression level of all annotated ecdysteroid genes were found to be higher in the females than males. Similar to sesquiterpenoid, the gene expression level of ecdysteroid genes of another bed bug species *C. lectularius* do not show any sex-biased expression (Supplementary Table S11; Supplementary Figure 26).

### MicroRNAs in *C. hemipterus* and other hemipterans

MicroRNAs are important post-transcriptional regulators in insects, but the microRNAs in *C. hemipterus* remain unexplored to date. Utilising the small RNA transcriptomes generated in this study, a total of 65 microRNAs could be confidently identified on 12 scaffolds (Figure 5A, Supplementary Table S14). Similar to other animals, some of these microRNAs were arranged in conserved genomic clusters, including miR-283/3477/12, miR-317/277, miR-750/1175, miR-9/306/79/9, miR-275/305, miR-87/87, let-7/miR-100, miR-263/263, and miR-71/2/13/13/2/2 (Figure 5A) (Chang et al 2016; Ikeda et al 2015; Marco et al 2013). By comparison of the microRNA expression levels between the two sexes in *C. hemipterus*, we also discovered that miR-12, miR-275, miR-279, miR-305, miR-306, miR-307 and miR-315 were expressed in a higher level in females than males, while miR-10, miR-1175, miR-1a, miR-219, miR-283, miR-3049, miR-316, miR-7, miR-87, miR-92, and miR-971 were in a higher expression level in males than females (Figure 5B, Supplementary Table S15, S16). How these microRNAs control the feeding behaviors between sexes remain to be elucidated.

**Figure 5.**
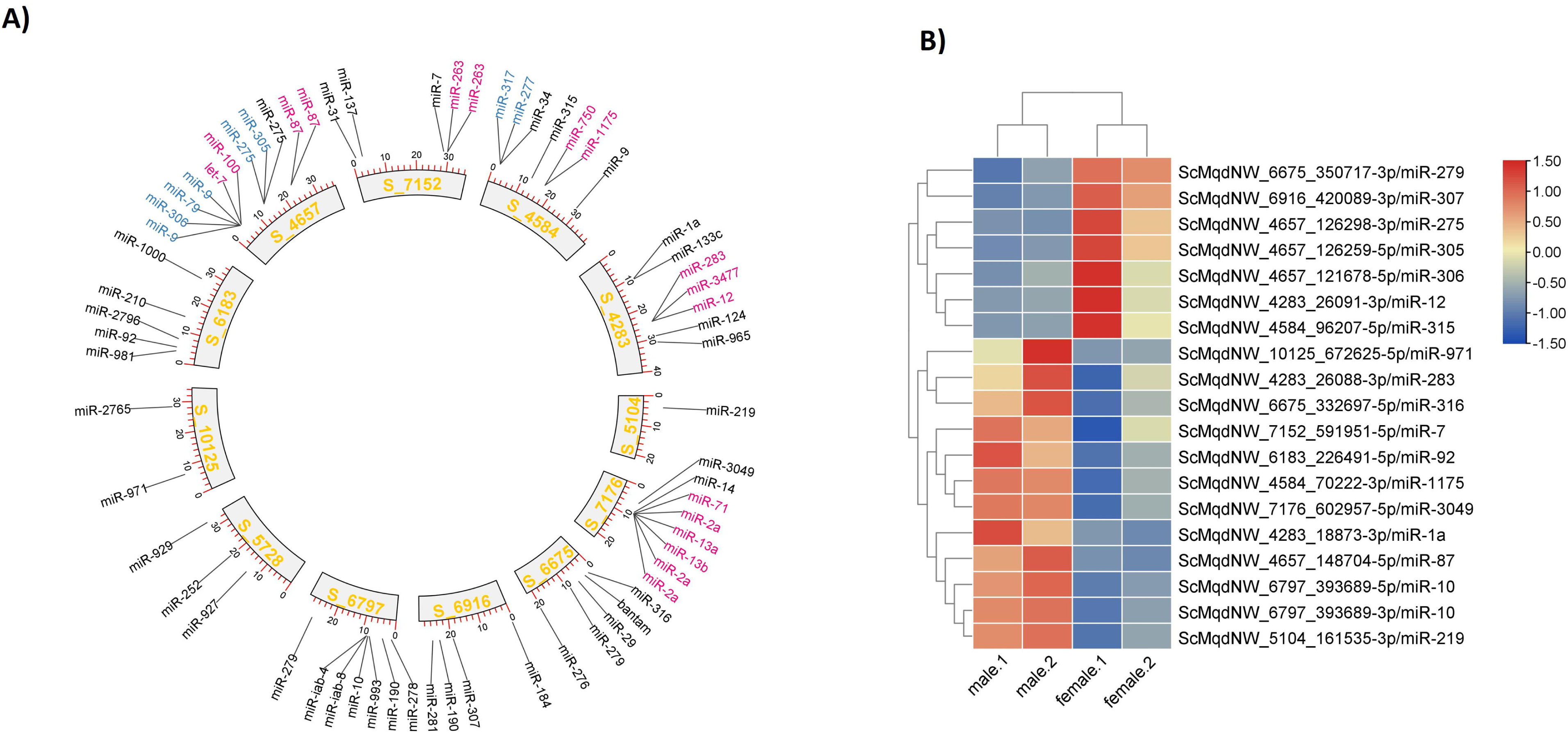
A) Genomic locations of microRNAs in tropical bed bug *Cimex hemipterus*. microRNA clusters are alternatively coloured in blue and red; B) MicroRNA differential expression genes.

## Conclusion

This study presents the first high-quality genome assembly for the tropical bed bug, which is scientifically important and has considerable medical relevance. Our work provides gene, transposable element, and microRNA annotations, and their detailed evolutionary analyses. More generally, our high-quality *C. hemipterus* genome, transcriptomic resources, and population data provides a useful reference point for further understanding of the biology, ecology, evolution, and infestation of insect pest.

## Data Availability

The final chromosome assembly was submitted to NCBI Assembly under accession number JASJUQ000000000 in NCBI. The raw reads generated in this study have been deposited to the NCBI database under the BioProject accessions: PRJNA713496, the genome annotation files were deposited in the Figshare (https://figshare.com/s/d9956fa56189ac8938c0).

## Abbreviations

BLAST: Basic Local Alignment Search Tool; BUSCO: Benchmarking Universal Single-Copy Orthologs; BWA: Burrows-Wheeler Aligner; cDNA: complementary DNA; gDNA: genomic DNA; kb: kilobase pairs; KEGG: Kyoto Encyclopedia of Genes and Genomes; LINE: long interspersed nuclear element; LTR: long terminal repeat; Mb: megabase pairs; NCBI: National Center for Biotechnology Information; PASA: Program to Assemble Spliced Alignments; SINE: short interspersed nuclear element; SRA: Sequence Read Archive.

## Supplementary Tables and Figures

Supplementary Table S1. Sequencing information of gDNA, mRNA, sRNA and population.

Supplementary Table S2. Statistics top 16 pseudomolecules of *C. hemipterus* genome.

Supplementary Table S3. Summary of BUSCO statistics.

Supplementary Table S4. Statistics on hemipteran genomes in the ncbi. Supplementary Table S5. Statistics on comparative genomes.

Supplementary Table S6. Hox and ParaHox genes in *C. lectularius* and *C. hemipterus*.

Supplementary Table S7. Expression of Hox and ParaHox genes in *C. hemipterus*.

Supplementary Table S8. Sesquiterpenoid genes of *C. hemipterus*.

Supplementary Table S9. Expression of sesquiterpenoid genes in *C. hemipterus* and *C. lectularius*.

Supplementary Table S10. Ecdysteroid genes of *C. hemipterus*.

Supplementary Table S11. Expression of ecdysteroid genes in *C. hemipterus* and *C. lectularius*.

Supplementary Table S12. Anticoagulation genes of *C. hemipterus*.

Supplementary Table S13. Expression of anticoagulation genes in *C. hemipterus*.

Supplementary Table S14. MicroRNA annotation.

Supplementary Table S15. MicroRNA expression table.

Supplementary Table S16. MicroRNA differential expression genes.

Supplementary Figure 1. Synteny between the X ,Y chromosome of the Heteroptera genomes (*Aelia acuminata*, GCA_911387785.2; *Acanthosoma haemorrhoidale* GCA_930367205.1; *Aradus depressus* GCA_963662175.1) downloaded from NCBI and the genome of this study.

Supplementary Figure 2. GenomeScope reports on the heterozygosity, repeat content and size of the genome with kmer 31

Supplementary Figure 3. Phylogenetic gene tree of apyrase genes identified in *C. hemipterus*

Supplementary Figure 4. Phylogenetic gene tree of salivary nitrophorin genes identified in *C. hemipterus*

Supplementary Figure 5. Phylogenetic gene tree of Kazal-type thrombin inhibitor genes identified in *C. hemipterus*

Supplementary Figure 6. Genomic location and expression profile of Far-17/AIG1 protein coding genes

Supplementary Figure 7. Classification of tetraspanin in *C. hemipterus* and other outgroup taxa

Supplementary Figure 8. Summary of tetraspanin in *C. hemipterus*

Supplementary Figure 9. Phylogenetic gene tree of Hox and ParaHox identified in *C. hemipterus* and *C. lectularius*

Supplementary Figure 10. Phylogenetic gene tree of ACAT identified in *C. hemipterus*

Supplementary Figure 11. Phylogenetic gene tree of HMGCS identified in *C. hemipterus*

Supplementary Figure 12. Phylogenetic gene tree of HMGCR identified in *C. hemipterus*

Supplementary Figure 13. Phylogenetic gene tree of MVK identified in *C. hemipterus*

Supplementary Figure 14. Phylogenetic gene tree of PMVK identified in *C. hemipterus*

Supplementary Figure 15. Phylogenetic gene tree of DPMD identified in *C. hemipterus*

Supplementary Figure 16. Phylogenetic gene tree of FPPS identified in *C. hemipterus*

Supplementary Figure 17. Phylogenetic gene tree of FNTB identified in *C. hemipterus*

Supplementary Figure 18. Phylogenetic gene tree of STE24 identified in *C. hemipterus*

Supplementary Figure 19. Phylogenetic gene tree of ICMT identified in *C. hemipterus*

Supplementary Figure 20. Phylogenetic gene tree of PCYOX1 identified in *C. hemipterus*

Supplementary Figure 21. Phylogenetic gene tree of ALDH3 identified in *C. hemipterus*

Supplementary Figure 22. Phylogenetic gene tree of JHAMT identified in *C. hemipterus*

Supplementary Figure 23. Phylogenetic gene tree of CYP15A1 identified in *C. hemipterus*

Supplementary Figure 24. Gene expression level of sesquiterpenoid genes in *C. hemipterus* and *C. lectularius*

Supplementary Figure 25. Phylogentic gene tree of ecdysteroid genes, Nvd, Spo/Spok, Phm, Dib, Sad, and Shd, identified in *C. hemipterus*

Supplementary Figure 26. Gene expression level of ecdysteroid genes in *C. hemipterus* and *C. lectularius*

